# Large donor CRISPR for whole-CDS replacement of cell adhesion molecule LRRTM2

**DOI:** 10.1101/2024.05.03.592283

**Authors:** Stephanie L Pollitt, Aaron D Levy, Michael C Anderson, Thomas A Blanpied

## Abstract

The cell adhesion molecule LRRTM2 is crucial for synapse development and function. However, our understanding of its endogenous trafficking has been limited due to difficulties in manipulating its coding sequence (CDS) using standard genome editing techniques. We instead replaced the whole LRRTM2 CDS by adapting the recent CRISPR method Targeted Knock-in using Two Guides (TKIT), enabling complete control of LRRTM2. In primary rat hippocampal cultures, N-terminally tagged, endogenous LRRTM2 was found in 80% of synapses, and synaptic LRRTM2 content correlated with PSD-95 and AMPAR levels. LRRTM2 was also enriched with AMPARs outside synapses, demonstrating the sensitivity of this method to detect relevant new biology. Finally, we leveraged total genomic control to increase the synaptic levels of LRRTM2 via simultaneous mutation of its C-terminal domain, which did not correspondingly increase AMPAR enrichment. The coding region of thousands of genes span lengths suitable for whole-CDS replacement, suggesting this simple approach will enable straightforward structure-function analysis in diverse cellular systems.

## Introduction

Synaptic organizing molecules, or cell adhesion molecules (CAMs), are unique transmembrane proteins that reach across the synaptic cleft, where they play critical roles in both cis-and transsynaptic biological processes from synaptogenesis to receptor trafficking and plasticity. The trafficking and expression of CAMs are tightly regulated, and mutations in CAM genes can severely disrupt neuronal function and are linked to neurological disease^1–3^. Accordingly, alterations of CAM expression levels, such as due to the use of exogenous expression for visualization and mutational analysis, can alter neuronal function and cloud interpretations of CAM biology. It is therefore necessary to measure and manipulate the endogenous protein itself to advance our understanding of CAMs. However, endogenous structure-function studies have been challenging, particularly because one critical approach is to mutate CAM sequences at multiple locations simultaneously. For example, N-terminal, extracellular tagging is ideal for most CAMs to visualize surface-expressed protein, but many critical protein interaction sites are quite distant from the N-terminus or even intracellular, as CAM C-tails play important roles in their anchoring and synaptic signaling. Standard genome editing techniques have largely been insufficient to make these changes simultaneously.

One example of a CAM for which new tools are needed is the synaptic CAM Leucine-Rich Repeat Transmembrane Protein 2 (LRRTM2). LRRTM2 resides in the post-synaptic membrane, where it binds to PSD-95 at its C-terminus and interacts trans-synaptically with presynaptic neurexins^4,5^. LRRTM2 exerts considerable power over synaptic function at mature synapses, largely as a result of its influence over AMPA receptor (AMPAR) anchoring and abundance at synapses. Knockdown of LRRTM2 affects basal AMPAR synaptic enrichment and synaptic strength^4^, and dual knockdown or knockout of LRRTM2 and its sister protein LRRTM1 abolishes long term potentiation (LTP) in hippocampal CA1 neurons ^6,7^. Furthermore, enzymatic cleavage of the LRRTM2 extracellular domain in a knockdown-rescue context disorganizes AMPAR subsynaptic distribution within minutes, and reduces total synaptic AMPAR content over the following hours^8^. LRRTM2 also plays a dose-dependent role in synaptogenesis, where over-or under-expression leads to corresponding changes in synapse density^4,9^. These results highlight the central role LRRTM2 plays in regulating synaptic strength at mature synapses and motivate detailed analysis of its molecular mechanisms.

As is true for many proteins, our understanding of LRRTM2 cellular functions has relied on genetic knockout and overexpression or antibody-based immunocytochemistry. Each of these approaches has considerable shortcomings. In particular, methods that manipulate LRRTM2 expression levels dramatically alter synaptic development and can engage compensatory mechanisms from related CAMs^4,9^. The use of knockdown-rescue can reduce overexpression phenotypes^4,8^, but expression via exogenous promoters can nevertheless cause diverse effects that are difficult to adequately monitor. While some studies have utilized antibody detection of endogenous LRRTM2, the low signal-to-noise ratio and lack of possibility to manipulate the protein sequence limits their scope^10,11^. Knock-in mouse models could bridge this gap; however LRRTM2 has numerous binding partners, and making a separate mouse model for each mutation is expensive and inefficient. Disappointingly, traditional CRISPR knock-in approaches to tag the LRRTM2 N-terminus have proven infeasible due to a lack of suitable PAM sites in the protein encoding region, and N-terminal knock-ins could threaten the integrity of the reading frame due to possible indels. C-terminal tagging is less than ideal due to inability to accurately identify protein present on the cell surface, and likely disrupts the PDZ-binding motif. Recently, the CRISPR approach TKIT (Targeted Knock-In with Two guides) was used to replace a protein-coding exon using two intron-localized guide RNA sequences^12^. This approach allowed N-terminal tagging of proteins without shifting the reading frame, as any Cas9-induced indels occur in intronic regions. We considered whether this strategy could be expanded and adapted to permit tagging and mutagenesis anywhere within a protein where the entire protein coding sequence (CDS) can be replaced. To test the feasibility of this approach while investigating LRRTM2 trafficking, we have applied whole-CDS replacement TKIT to the rat *Lrrtm2* gene.

In this work, we demonstrate whole-CDS replacement in neurons as well as the simultaneous tagging and mutagenesis of the *Lrrtm2* coding region. We utilized endogenously tagged LRRTM2 to measure its synaptic enrichment as well as its relationship with AMPAR enrichment at synapses. To enable rapid identification and measurement of knock-in cells, we also developed a strategy to express a marker conditional on successful knock-in, enabling new and flexible experimental designs. Finally, we demonstrated the power of this technique by combining N-terminal tagging with distal point mutations to increase synaptic surface enrichment of LRRTM2. Together, our findings provide context for our understanding of LRRTM2 function as well as a novel method for future structure-function studies in neurons and other post-mitotic cells.

## Results

### Two-guide CRISPR approach TKIT can be adapted for whole-CDS replacement of LRRTM2

To decipher the trafficking patterns of the key synaptic cell adhesion molecule LRRTM2, we designed an approach to simultaneously measure and manipulate the protein while maintaining its endogenous transcriptional regulation and expression level. HITI is a common CRISPR genome editing method used in post-mitotic cells such as neurons that allows for DNA insertions that could be useful for tagging native LRRTM2^13^. However, HITI is limited to insertion at single sites, which for LRRTM2 would only allow either an N-terminal tag or protein mutagenesis, not both. In addition, HITI targets PAMs to make double stranded breaks within the coding region, and therefore may generate indels that alter the reading frame unpredictably (Figure 1A, left). Therefore, we considered whether we could edit the entire mature LRRTM2 CDS by adapting the exon-replacement technique Targeted Knock-In with Two guides (TKIT)^12^. Instead of targeting a single PAM site within the coding region, TKIT utilizes two guide RNAs targeting PAMs in the non-coding regions flanking an exon, which directs Cas9 to excise the exon entirely and enables its replacement with a synthetic exon (Figure 1A, middle). This has the advantage of improving knock-in efficiency since multiple guide RNAs and PAM sites can be tested throughout the intronic sequences, and it reduces the effects of indels as they will occur outside the coding region. TKIT has previously been used to replace the 5’-most coding exon of diverse neurotransmitter receptors with a synthetic version that includes an N-terminal fluorescent tag^12^. We wondered whether this approach could be modified such that the entire coding region of a gene, in our case LRRTM2, could be replaced instead (Figure 1A, right). Notably, nearly the entire LRRTM2 protein is encoded in a single exon, with the 5’ UTR and only 4 bases of signal peptide in exon 1, and the rest of the coding region, plus the 3’ UTR, in exon 2 (Figure 1B). This genomic organization offered the attractive possibility of replacing the entire protein-coding sequence of LRRTM2 with an entirely customizable one using TKIT. Therefore, we modified the TKIT approach to replace much of the second exon, including all the coding sequence contained therein. Given that the signal peptide is cleaved co-translationally, this approach would allow us to replace the entire mature LRRTM2 protein with a donor sequence of our choosing.

**Figure 1:**
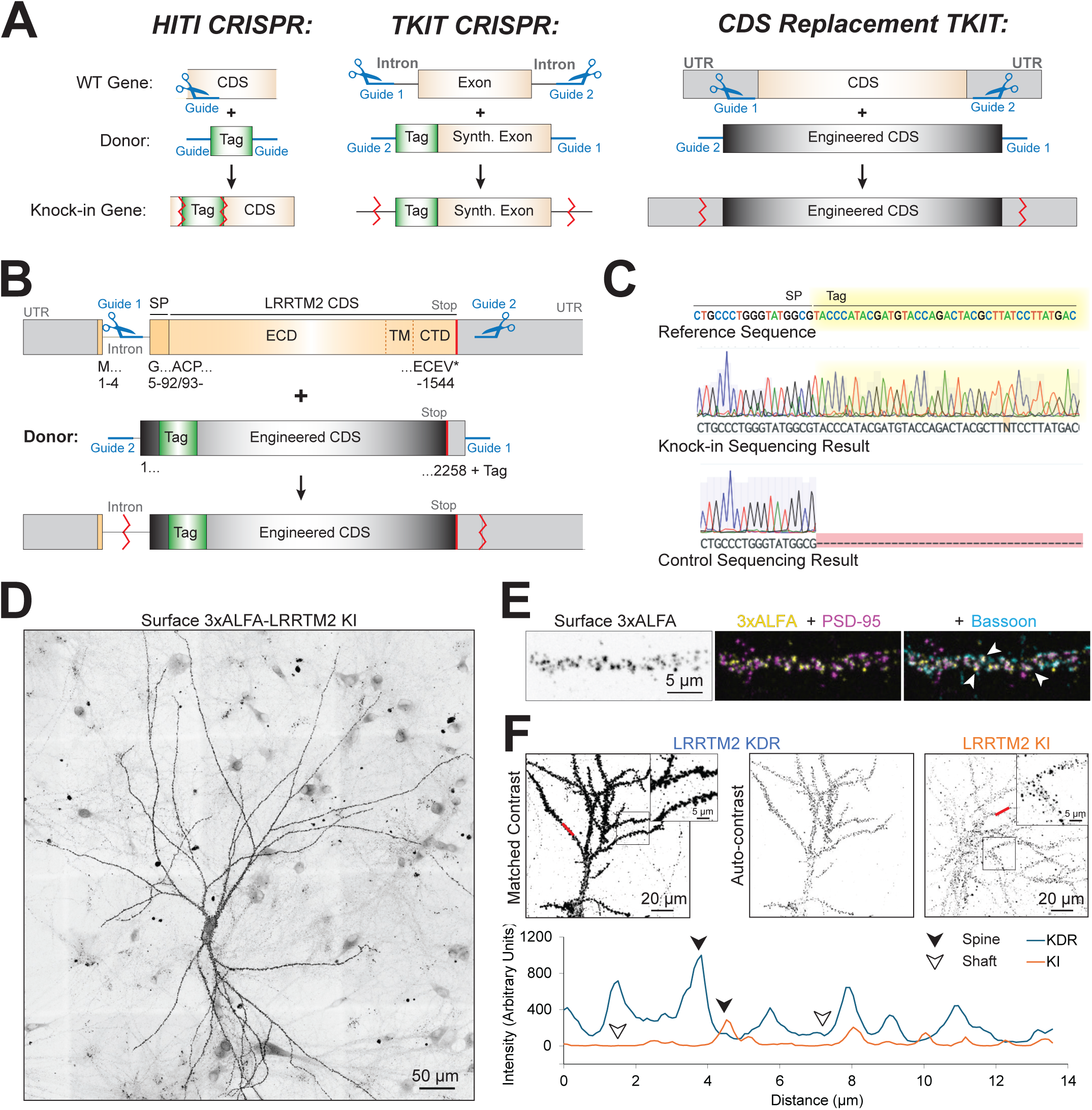
Successful whole-CDS replacement of LRRTM2 **A)** Diagram of traditional HITI, TKIT, and whole-CDS replacement CRISPR methods. Locations of possible Cas9-induced indels are indicated with red zigzag lines. **B)** Diagram of the *Rattus norvegicus Lrrtm2* gene with CRISPR guide locations. Note that the entire coding sequence (orange), minus 4 bases of the signal peptide that is cleaved from the mature protein, is confined to exon 2. Guide 1 anneals to the intronic region and guide 2 to the 3’UTR. The replacement donor (Engineered CDS, black) contains an N-terminal epitope tag between the signal peptide and mature protein. **C)** Genomic sequencing shows positive knock-in result. Samples of genomic DNA from either the guides/donor virus alone (Control) or the guides/donor and Cas9 viruses (Knock-in, KI) underwent two rounds of PCR amplification, first to amplify the *Lrrtm2* gene, then again to amplify the knock-in tag (no product in the control condition). Sanger sequencing shows the 3xHA tag sequence in the KI condition, without indels. **D)** Image of a cultured hippocampal knock-in neuron, stained for 3xALFA-LRRTM2. Sparse knock-in permits identification of synaptic puncta along knock-in vs neighboring dendrites. **E)** Images of a dendrite from a knock-in neuron, stained for 3xALFA-LRRTM2 (yellow), PSD-95 (magenta), and pre-synaptic Bassoon (cyan). LRRTM2 puncta are largely synaptic (white arrows). **F)** Endogenous LRRTM2 (KI, right) is expressed at far lower levels than typical exogenous expression via knockdown-rescue (KDR, left). Matched staining, imaging, and look-up tables show that while both KDR and KI show synaptic localization, knockdown-rescue ALFA labeling is considerably higher than knock-in both at synapses and between (blue and orange, respectively, line scan, lower right). Dark arrows indicate spine enrichment, white arrows dendritic signal. Line scan locations shown with red lines in the respective images.

To replace the LRRTM2 coding sequence using TKIT, we designed a 5’ guide RNA (guide 1) targeting a PAM site within intron 1 of *Rattus norvegicus Lrrtm2* and a 3’ guide RNA (guide 2) targeting a PAM site after the *Lrrtm2* stop codon within the 3’UTR, resulting in a large span of replacement that exceeds those of previous efforts^12,14,15^. The replacement donor sequence was engineered using rat genomic DNA and subcloned to include an epitope tag between the signal peptide and LRRTM2 N-terminal regions. The donor was also flanked by flipped and reversed guide sequences to allow reversed integration to be fixed, as used in both HITI and TKIT^12,13^ (Figure 1B). We tested several N-terminal tags for both specific labeling intensity and non-specific background from the affinity reagent using exogenous LRRTM2 expression in HEK cells (Supplemental Figure 1). We found that the ALFA nanobody^16^ displayed lower non-specific signal than an HA antibody plus secondary antibody. Further, we were able to increase specific staining of the ALFA epitope by adding a triple ALFA tag without interstitial linkers, which had approximately 50% higher fluorescence intensity than either the single ALFA tag or a triple ALFA tag with linkers. Therefore, we utilized a triple ALFA tag without linkers in the donor sequence for fluorescent labeling (note that for genomic PCR and sequencing experiments, a triple HA tag was used instead for convenience). As transfection efficiency of cultured neurons is low, we delivered the necessary DNA with two lentiviruses, one to express the guides and donor sequence and a second to express Cas9.

We first validated the success of LRRTM2 whole-CDS replacement by infecting dissociated embryonic rat hippocampal neurons with either Cas9 and LRRTM2 guides/donor lentiviruses or with guides/donor virus only as a control, then sequenced genomic DNA after three weeks in culture. Briefly, we utilized PCR to isolate the *Lrrtm2* gene and amplified knock-in-positive alleles using sequential PCRs (Figure 1B, bottom) and sequenced the resulting amplicons via Sanger sequencing. Positive sequences for the 3xHA tag were identified in the knock-in condition (Figure 1C, middle) but not in the control condition lacking Cas9 (Figure 1C, bottom), indicating that the tag was successfully incorporated into the correct gene locus and that knock-in was successful. Note that we did not introduce indels at the signal peptide-tag junction or at the tag-protein junction, as this region was contained entirely within the engineered donor sequence as is true for TKIT. We next evaluated whether the knocked-in 3xALFA tag would allow us to visualize the endogenous expression of LRRTM2. We performed surface fluorescent labeling of 3xALFA-LRRTM2 on live, dissociated hippocampal cultured neurons infected three weeks prior with the knock-in constructs, then fixed the cells and additionally immunolabeled for PSD-95 and the presynaptic scaffold protein Bassoon to visualize synapses. Fluorescent confocal imaging showed isolated cells with anti-ALFA labeling in discrete puncta along the cell body and dendrites (Figure 1D). Close inspection revealed these puncta typically occurred at synapses, as indicated by colocalization with both Bassoon and PSD-95, consistent with previous studies of LRRTM2^4,10^ (Figure 1E, especially arrowheads). In line with previous TKIT-based knock-ins, we commonly observed 20 to 30 ALFA-stained cells per coverslip with this method; note that this may be an underestimate of the knock-in efficiency as some cells may be knocked-in but not express LRRTM2, and some cells may not be co-infected by both viruses. These results together demonstrate that our knock-in was successful and could be used to visualize endogenous LRRTM2 in cultured neurons.

The previous best practice for visualizing LRRTM2 has been knockdown-rescue (KDR), which is frequently assumed to minimize overexpression due to the knockdown of endogenous expression. However, this method still typically relies on exogenous or unregulated promoters, which often express proteins at higher than endogenous levels. To compare knock-in LRRTM2 expression levels with those following KDR, we performed live surface ALFA labeling on cells infected with knock-in constructs and on cells transfected with a knockdown-rescue 3xALFA-LRRTM2 construct. As expected, ALFA staining was clearly far higher in the KDR condition than the knock-in (Figure 1F). Further, while the expression pattern of both was largely synaptic, LRRTM2 signal in the KDR condition showed higher and more consistent levels of LRRTM2 at each spine (Figure 1F, line scan, dark arrows), and also showed higher ALFA labeling along the dendrite outside of dendritic spines (Figure 1F, line scan, white arrows). This indicates that overexpression of LRRTM2, even following knockdown, can alter its surface and subcellular trafficking, highlighting the importance of labeling and measuring endogenous protein.

### Rapid identification of whole-CDS knock-in cells

As is true for many endogenously labeled proteins, the low endogenous expression levels of LRRTM2 made visualizing knock-in cells or tracing their morphology for analysis difficult. We therefore leveraged our whole-CDS replacement system to design a knock-in conditional marker. The conditional effect was achieved by adding an Internal Ribosome Entry Sequence (IRES) and Cre gene after the stop codon in the LRRTM2 donor containing the 3xALFA tag. This donor could allow for expression of a FLExible marker protein^17^ to be conditional upon successful knock-in of 3xALFA-LRRTM2 (Figure 2A). To test the utility of this approach, we infected neurons with three lentiviruses expressing the 3xALFA-LRRTM2-IRES-Cre donor and guide RNAs, Cas9, and FLEx-mTagBFP2, expecting that when the 3xALFA-LRRTM2-IRES-Cre donor is knocked into the *Lrrtm2* locus and the mRNA translated, Cre will also be expressed and able to induce FLEx-mTagBFP2 expression (Figure 2A, bottom). This strategy successfully produced cells expressing both knock-in 3xALFA-LRRTM2 and knock-in-dependent mTagBFP2 cell fill, facilitating rapid visual identification on the microscope (Figure 2B). Notably, the 3xALFA tag and IRES-Cre sequences are 1449 bp away from one another on the donor sequence, a simultaneous knock-in impossible to make by other methods. The donor used was also 3925 bp long, over two times larger than the largest TKIT donor previously reported^12^ and among the largest we are aware of successfully being knocked in using NHEJ-based CRISPR methods. We observed a fraction of neurons that were positive for mTagBFP2 without detectable surface ALFA labeling (16%), presumably reflecting cell types with successful knock-in that do not express or only transiently expressed LRRTM2. Cells in this category were excluded from further analysis. A small number of neurons (8%) were positive for surface ALFA staining without detectible mTagBFP2 expression, potentially due to a lack of coinfection by all three viruses. Nevertheless, expression of the marker protein was routinely high enough to rapidly identify presumed knock-in cells for image acquisition. An important advantage of this approach is the potential to use alternative conditional reporters. To illustrate this, we utilized a FLEx-IRES-EGFP lentivirus in place of the FLEx-mTagBFP2 and observed EGFP-positive knock-in cells (Figure 2C). Conceivably, any FLEx fluorescent protein, sensor, or optogenetic tool could be used to accommodate numerous experimental approaches, demonstrating the utility of our whole-CDS control.

**Figure 2:**
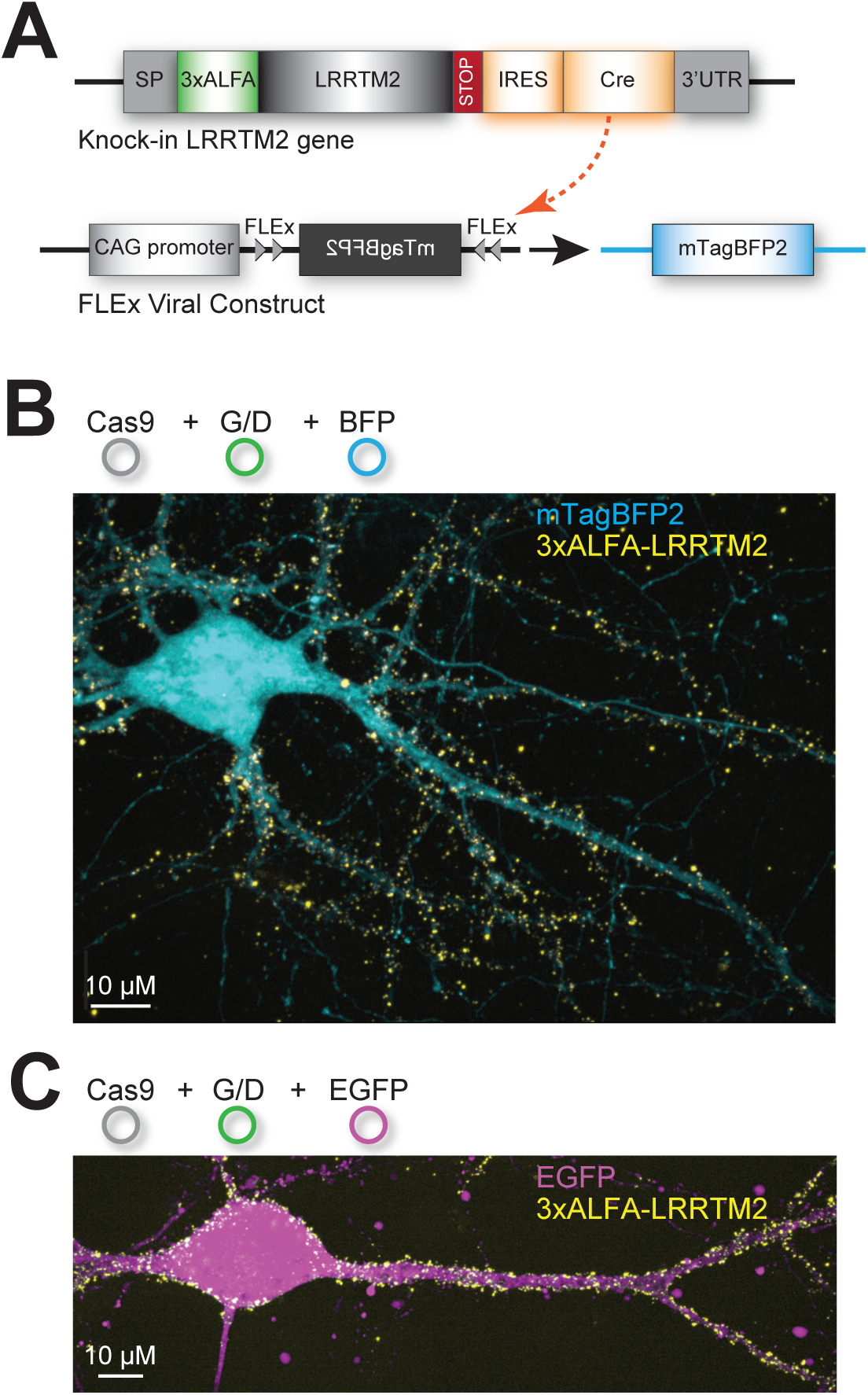
Rapid identification of whole-CDS knock-in cells **A)** Schematic of knock-in conditional marker implementation. An Internal Ribosome Entry Site (IRES) sequence and a Cre gene were inserted into the donor construct between the LRRTM2 STOP codon and the 3’UTR (IRES-Cre, orange). Thus, Cre is expressed once the donor is successfully incorporated into the LRRTM2 gene. A separate lentiviral construct was used to deliver FLEx-mTagBFP2 cell fill, which can be targeted by the knocked-in Cre. **B)** Knock-in neuron expresses the mTagBFP2 cell fill. Surface immunostained 3xALFA-LRRTM2 (yellow), cytosolic mTagBFP2 (cyan). **C)** The IRES-Cre approach permits flexibility in the conditional marker. Due to the flexible multi-virus approach, it is simple to swap mTagBFP2 for any other FLEx protein marker, such as EGFP (magenta). Three viral constructs were used: Cas9, Guides/Donor, and FLEx marker.

### Endogenous LRRTM2 labeling enables investigation of LRRTM2 trafficking and synaptic enrichment

We next set out to deploy our LRRTM2 knock-in and conditional reporter system to characterize LRRTM2 expression and synaptic trafficking at endogenous levels. We first quantified the average synaptic LRRTM2 expression level per cell and found a wide range of intensities across cells (Figure 3A). This suggests that LRRTM2 expression varies within the transcriptomic profile of neurons in these CA1-enriched hippocampal cultures, indicating possible cell-intrinsic control of LRRTM2 expression. We considered that some of the cells with higher LRRTM2 intensity might be homozygous for the 3xALFA-LRRTM2 knock-in, and those with lower expression might be heterozygous. However, we observed a smooth, rather than bimodal, distribution of average synaptic intensity (Figure 3B), consistent with regulation of LRRTM2 expression levels by activity history or other cell-specific mechanism.

**Figure 3:**
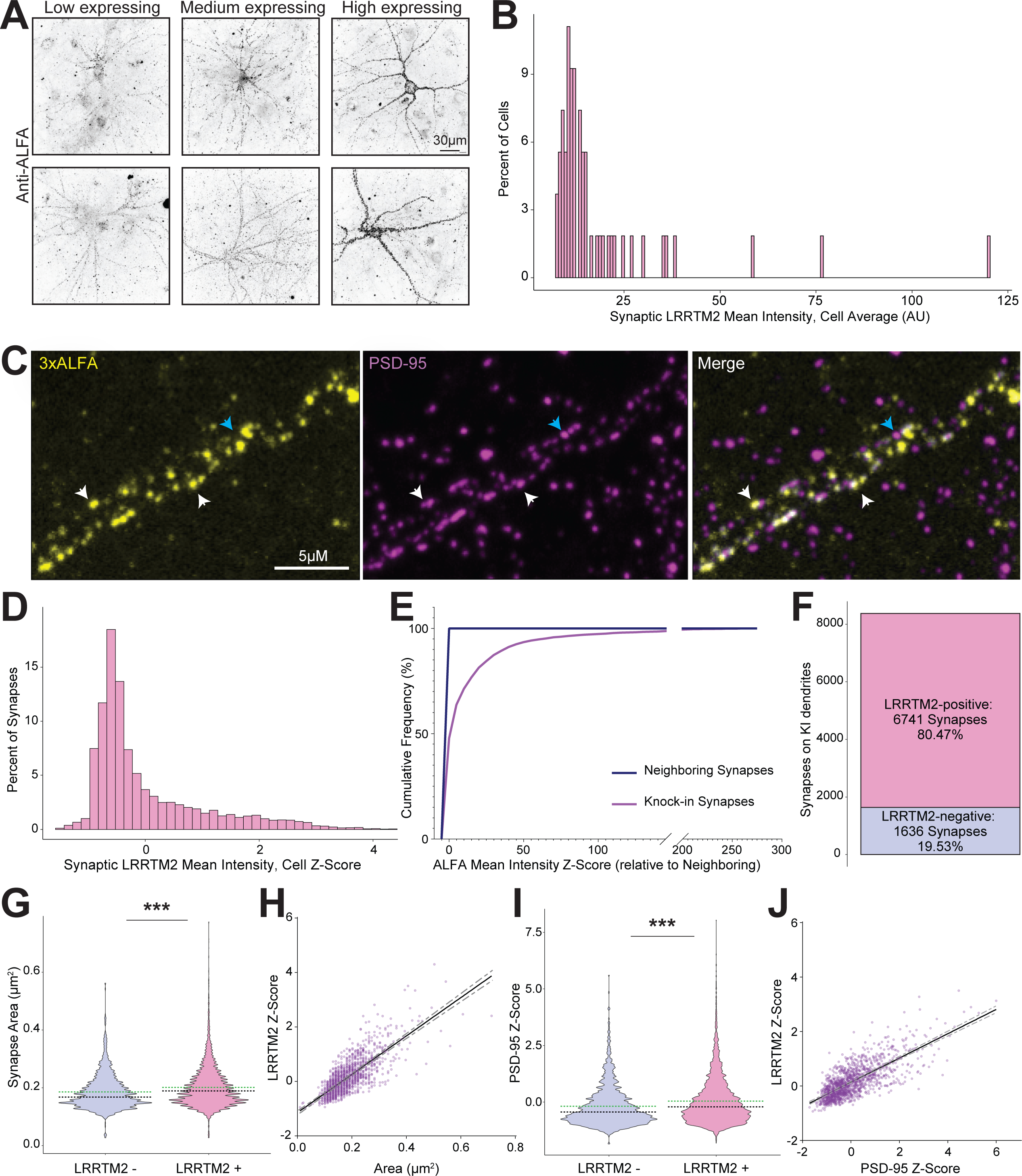
Variation in endogenous synaptic LRRTM2 content reflects key markers of synaptic strength. **A)** Exemplar hippocampal neurons expressing 3xALFA-LRRTM2 at low, medium, and high levels. **B)** Histogram of synaptic LRRTM2 expression across the dendrite, averaged by cell. **C)** Image of an example stretch of dendrite expressing knock-in 3xALFA-LRRTM2 (yellow) immunolabeled for PSD-95 (magenta). LRRTM2 is highly synaptic in localization (exemplars, white arrows), but does not appear in every synapse on the dendrite (blue arrow). **D)** Distribution of synaptic LRRTM2 mean intensity along knock-in dendrites, normalized for cellular variability using a per-cell Z-scoring method. **E)** Cumulative frequency plot of ALFA mean intensity at synapses in knock-in neurons (pink) or neighboring non-knock-in dendrites (blue). Z-scores for LRRTM2 mean intensity measures were calculated from the mean and standard deviation of synaptic ALFA signal within neighboring non-knock-in synapses. **F)** Bar plot of synapses along knock-in dendrites categorized as containing (pink) or lacking (blue) LRRTM2. The 95^th^ percentile of non-knock-in mean intensity Z-scores was used to delineate the cutoff between LRRTM2-containing and -lacking synapses. **G)** Violin plot comparing synapse area along knock-in dendrites that do or do not contain LRRTM2 (pink or blue, respectively). (p < 2.2^-16^) **H)** Scatter plot correlating synapse area with synaptic LRRTM2 content (raw integrated density Z-score) in LRRTM2-containing synapses. **I)** Violin plot comparing PSD-95 content within synapses along the knock-in dendrites that do or do not contain LRRTM2. (p < 2.2^-16^) **J)** Scatter plot of PSD-95 content with synaptic LRRTM2 content at LRRTM2-containing synapses. In violin plots, green lines represent means and black lines represent medians for each distribution. Scatter plots have been averaged across every 5 ranked data points for visibility; statistics are calculated from raw data. Lines show linear regression with gray dotted lines representing 95% confidence intervals.

LRRTM2 is synaptically localized and plays a key role in synaptic function^4,6,7^. However, our observation that LRRTM2 puncta were more discrete and variable with our knock-in than with KDR due to overexpression artifacts (Figure 1F) indicated its endogenous distribution may be more nuanced. We therefore investigated in detail the variability of endogenous LRRTM2 content at individual synapses in 3xALFA-LRRTM2 knock-in neurons, immunolabelled for PSD-95 with a fluorescently labeled nanobody (Figure 3C). Despite prominent colocalization at synapses (Figure 3C, white arrows), we observed that the quantity of LRRTM2 varied substantially between synapses (coefficient of variation 1.514) and that, in fact, not every synapse along dendrites from knock-in cells contained LRRTM2 (Figure 3C, blue arrow). To better understand the synaptic variation of LRRTM2 content while accounting for cellular variability of LRRTM2 expression levels (Figure 3A), we measured LRRTM2 intensity within synapses and normalized for variable cellular expression by calculating cell-based intensity Z-scores for each synapse. We identified synaptic puncta as PSD-95 regions of interest (ROIs) using a semi-automated synapse detection analysis^18,19^ of images from 54 knock-in neurons across 3 culture replicates and calculated Z-scores for each synapse. We measured the area as well as the fluorescence intensity of each labeled protein within these synapse ROIs, and normalized for their variable cellular expression by calculating cell-based intensity Z-scores for each synapse using the cellular average and standard deviation for LRRTM2 or PSD-95, respectively (Figure 3D). This analysis clearly showed that synaptic LRRTM2 content varied considerably within individual neurons, even when normalized for cellular variability.

To define a synapse as lacking LRRTM2, we sought to establish a baseline by examining the off-target ALFA staining. This was done by measuring ALFA intensity at neighboring synapses from non-knock-in neurons, reflecting non-specific staining and noise, which as expected had Z-scores near 0 (Figure 3E, Neighboring Synapses). To compare these two distributions, we recalculated the Z-scores of knock-in LRRTM2 intensity using the mean and standard deviation of ALFA signal in neighboring PSDs, and found the synapses along the knock-in dendrite were, as expected, far brighter in ALFA signal than the neighboring background staining. (Figure 3E). We then used the 95^th^ percentile of non-knock-in synapse ALFA staining z-scores to delineate a cutoff between LRRTM2-containing and LRRTM2-lacking synapses within the knock-in dataset. Using this cutoff, approximately 80% of the synapses along knock-in dendrites contained LRRTM2 (Figure 3F). This is slightly higher than a previous estimate utilizing LRRTM2 antibody staining in DIV13-15 mouse hippocampal cultures^11^, which could represent a developmental or species difference, but also could be a reflection of our ability to more precisely estimate background noise from cells that unequivocally lacked the knock-in and therefore accurately identify true signal.

Our observation of a synapse subtype lacking LRRTM2 raises additional questions about what other differences these synapses exhibit relative to their LRRTM2-containing neighbors. Given that LRRTM2 plays a role in synaptic plasticity and binds to PSD-95, one explanation of its synaptic variability is that synapses with higher LRRTM2 levels might be larger and contain more PSD-95. Indeed, we found that synapses with LRRTM2 exhibited larger PSDs than those without (Wilcox test: p < 2.2e-16, r-statistic: -0.142; Figure 3G). Further, within the population of LRRTM2-containing synapses, larger synapses contained more LRRTM2 protein (slope: 7.061, R^2^: 0.226; Figure 3H). Our interpretation is further supported by examining PSD-95 content – synapses with LRRTM2 contained more PSD-95 than those without LRRTM2 (Wilcox: p < 2.2e-16, r-statistic: -0.149; Figure 3I), and PSD-95 and LRRTM2 intensities were positively correlated (Slope: 0.442, R-squared: 0.183; Figure 3J). These data indicate that the subcellular trafficking of LRRTM2 to synapses is correlated with both synapse size and the amount of PSD-95 present. While previous studies have indicated that overexpressed LRRTM2 does not require PSD-95 binding to be enriched at synapses^10^, our data indicates that nonetheless LRRTM2 trafficking scales with these key markers of synaptic strength.

### Synaptic AMPA receptor content scales with LRRTM2 content

Previous studies utilizing LRRTM1/2 KDR or LRRTM1/2 KO replacement with LRRTM2 have established that LRRTM2 stabilizes AMPARs in synapses at baseline and anchors new AMPARs after long-term potentiation^6,7^. Despite these findings, it has been difficult to disentangle how AMPAR trafficking compares with endogenous expression levels of LRRTM2. Furthermore, our ability to visualize endogenous LRRTM2 and identify synapses lacking it gives us the opportunity to compare AMPAR content between these two groups of synapses. We hypothesized that synapses lacking endogenous LRRTM2 would also have reduced AMPAR content relative to synapses with LRRTM2. To examine the relationship between the synaptic enrichment of LRRTM2 and AMPARs, we immunolabeled surface AMPA receptors simultaneously with ALFA-LRRTM2, fixed and stained the neurons for PSD-95 to identify synaptic puncta, and measured synaptic LRRTM2 and AMPAR intensity from the same synapses (Figure 4A). As in Figure 3, we normalized both the LRRTM2 and AMPAR raw integrated density signals using Z-scores based on respective means in each cell, then assessed synapses along knock-in dendrites for the presence or absence of LRRTM2. Consistent with the role of LRRTM2 in AMPAR anchoring, LRRTM2-lacking synapses contained less surface AMPAR signal than synapses with LRRTM2 (Wilcox: p < 2.2e-16, r-statistic: -0.136; Figure 4B). While LRRTM2-lacking synapses still exhibited AMPAR staining, this is likely due to stabilization by other mechanisms. When LRRTM2 was present, synapses with more LRRTM2 also had more AMPAR content on the surface (Slope: 0.3513, R-squared 0.1326; Figure 4C). While these results do not support a model where endogenous LRRTM2 is solely responsible for synaptic AMPAR anchoring, the correlation between AMPAR content and LRRTM2 content supports the hypothesis that LRRTM2 plays a strong role in control of AMPAR trafficking^6,7^.

**Figure 4:**
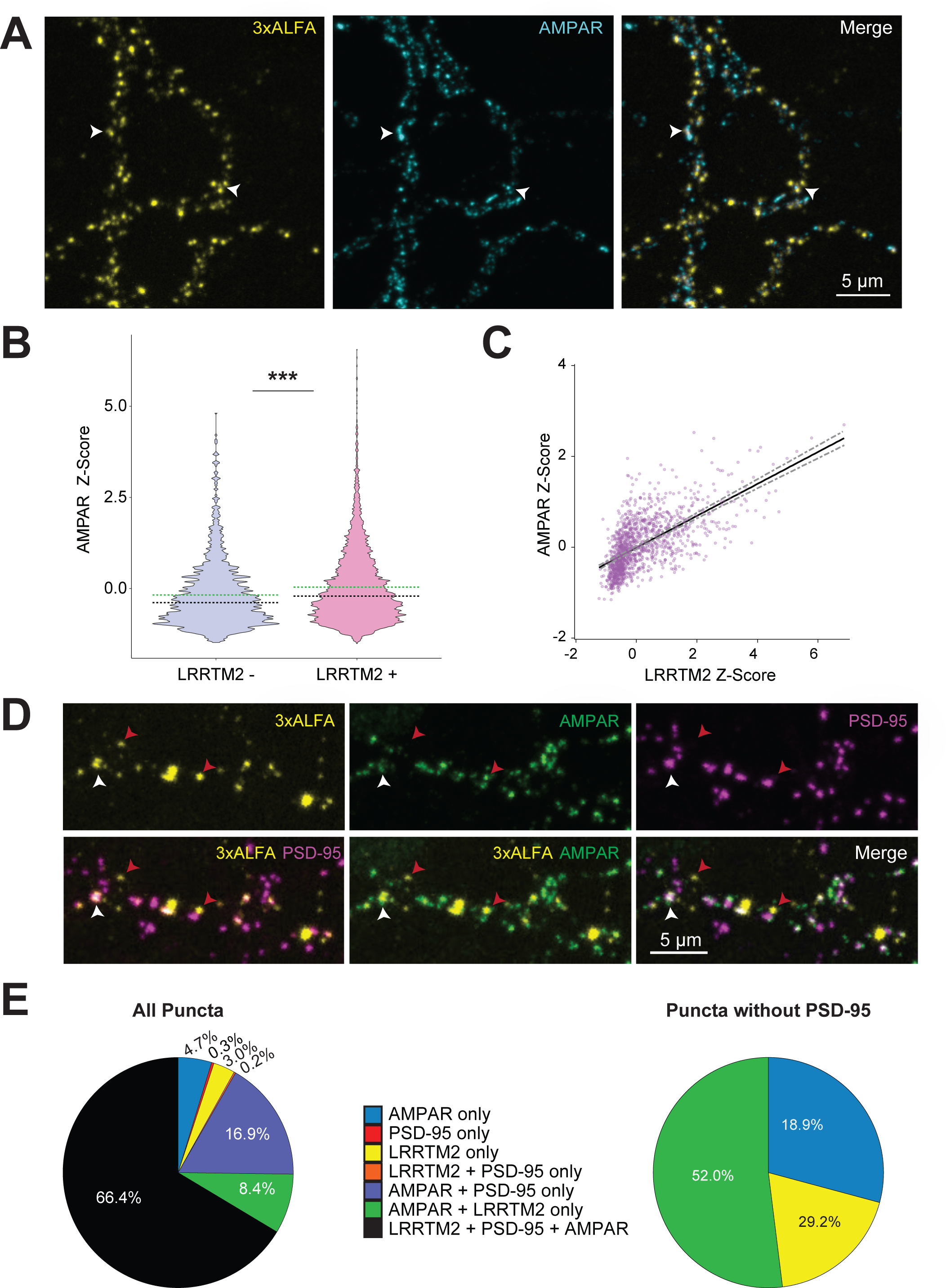
Synaptic AMPAR content scales with LRRTM2 content **A)** Images of a knock-in neuron labeled for 3xALFA-LRRTM2 (yellow) and surface AMPARs (cyan). Arrows show high co-enrichment of AMPARs with LRRTM2. **B)** Violin plot comparing synaptic surface AMPAR content along the knock-in dendrites that do or do not contain LRRTM2 (pink and blue, respectively). p < 2.2^-16^ Green lines represent means and black lines represent medians for each distribution. **C)** Scatter plot correlating AMPAR content (Raw Integrated Density z-score) with synaptic LRRTM2 content (Raw Integrated Density z-score) at LRRTM2-containing synapses. Scatter plot has been averaged across every 5 ranked data points for visibility; statistics are calculated from raw data. Lines show linear regression with gray dotted lines representing 95% confidence intervals. **D)** Images of triply labeled neurons with 3xALFA-LRRTM2 (yellow), PSD-95 (magenta), and surface AMPARs (green). Arrows indicate LRRTM2 clusters that do (white) or do not (red) co-enrich with PSD-95. **D)** Left: Pie chart depicting the proportions of LRRTM2, AMPAR, and/or PSD-95-containing puncta along knock-in dendrites. Right: Subset of left, AMPAR and/or LRRTM2 clusters without PSD-95 enrichment. Notably, LRRTM2 puncta outside of PSD-95 often also contain AMPAR clusters.

Surprisingly, close examination of these triply-labeled knock-in cells revealed occasional LRRTM2 puncta that fell outside of PSD-95-demarcated synapses (Figure 4D, red vs white arrows). It is well documented that extrasynaptic AMPAR pools play a role in regulating synapse strength, particularly during long-term potentiation^20,21^. Given the role of LRRTM2 in LTP and AMPAR stabilization, we therefore wondered whether these LRRTM2 puncta outside of PSD-95 might also contain AMPARs. To address this, we selected isolated dendrites from 23 knock-in cells across three culture replicates and identified puncta in each channel (surface AMPARs, PSD-95, and 3xALFA-LRRTM2) as ROIs for measurement. As the semi-automated detection was trained to detect puncta that are sized and shaped like synapses and ignore nonsynaptic labeling, these ROIs were hand-selected. We then measured the raw integrated density within these ROIs in all three channels, and qualitatively labeled each as positive or negative for the other labeled proteins based on its intensity, where intensities above the first local minimum of the distribution were described as “positive” for that corresponding protein (Figure 4E, left). Consistent with our previous automated analysis, when the hand-selected PSD-95 puncta were pooled together, we found similar numbers of LRRTM2-positive and - negative synapses (by hand: 79.7% positive and 20.2% negative; automated: 80.5% positive and 19.5% negative), validating our approaches. We then broke down the non-PSD-95-containing puncta into groups based on their protein expression (Figure 4E, right). Most such puncta contained both LRRTM2 and AMPARs together (52%), more than the puncta with either protein on its own (29% AMPAR only and 19% LRRTM2 only). The existence of LRRTM2/AMPAR puncta outside PSD-95 synapses could be partially explained to be synapses that do not contain PSD-95 and instead have another scaffold such as SAP-102 or PSD-93^36,37^. However, given that such synapses are relatively rare in mature hippocampal neurons, it’s likely that at least some of them are genuinely extrasynaptic. This raises the exciting possibility that along with its known localization within synapses, LRRTM2 could co-diffuse with extrasynaptic AMPARs, and possibly play a role in their trafficking there.

### Simultaneous mutagenesis and tagging enabled by whole-CDS replacement permits analysis of mutation-induced changes in LRRTM2 surface expression

A major advantage of whole-CDS replacement is the potential it offers to easily edit a protein of interest at multiple sequence points, as we have already demonstrated by adding IRES-Cre at the C-terminus simultaneously with the N-terminal 3xALFA tag. To demonstrate the power of whole-CDS replacement of LRRTM2, we modified our synthetic donor sequence to include, along with the IRES-Cre, a functional mutation near the C-terminus expected to alter LRRTM2 trafficking. Previous work has shown that surface enrichment of exogenously expressed LRRTM2 can be manipulated through two point mutations in the C-terminal domain, Y501A and C504A^22,23^, which are hypothesized to alter the membrane-targeting trafficking mechanisms of LRRTM2^23^. Termed “YACA”, these mutations induce a 400% increase in surface LRRTM2 levels in a knockdown-rescue context^22^; however as we established, the KDR protein trafficking pattern may not be the same as endogenous. Our whole-CDS replacement model offers the ideal platform on which to test this. We introduced the YACA mutations into our donor construct to create a 3xALFA-LRRTM2-YACA-IRES-Cre knock-in (Figure 5A), infected neurons with lentivirus to generate wild type or YACA knock-ins, then measured AMPAR and LRRTM2 content in PSD-95-labeled synapses as above (54 wild type and 64 YACA neurons across 3 culture replicates; Figure 5B). On a per cell basis, average synaptic LRRTM2-YACA content was 24.6% higher than wildtype (Wilcox: p < 0.005, r-statistic = 0.280; Figure 5C). When we examined the per synapse distribution of LRRTM2 intensities between the two conditions, we found a similar trend where the distribution of YACA-containing synapses was right-shifted relative to wildtype (Wilcox: p < 2.2e-16; r-statistic = 0.102; Figure 5D). Therefore, while the YACA mutations did increase endogenous LRRTM2 surface trafficking as expected, the percent change was dramatically smaller than that observed in a KDR model. These findings reinforce the value of our whole-CDS replacement approach and demonstrate the power of simultaneous genomic editing at multiple sites within a protein.

**Figure 5:**
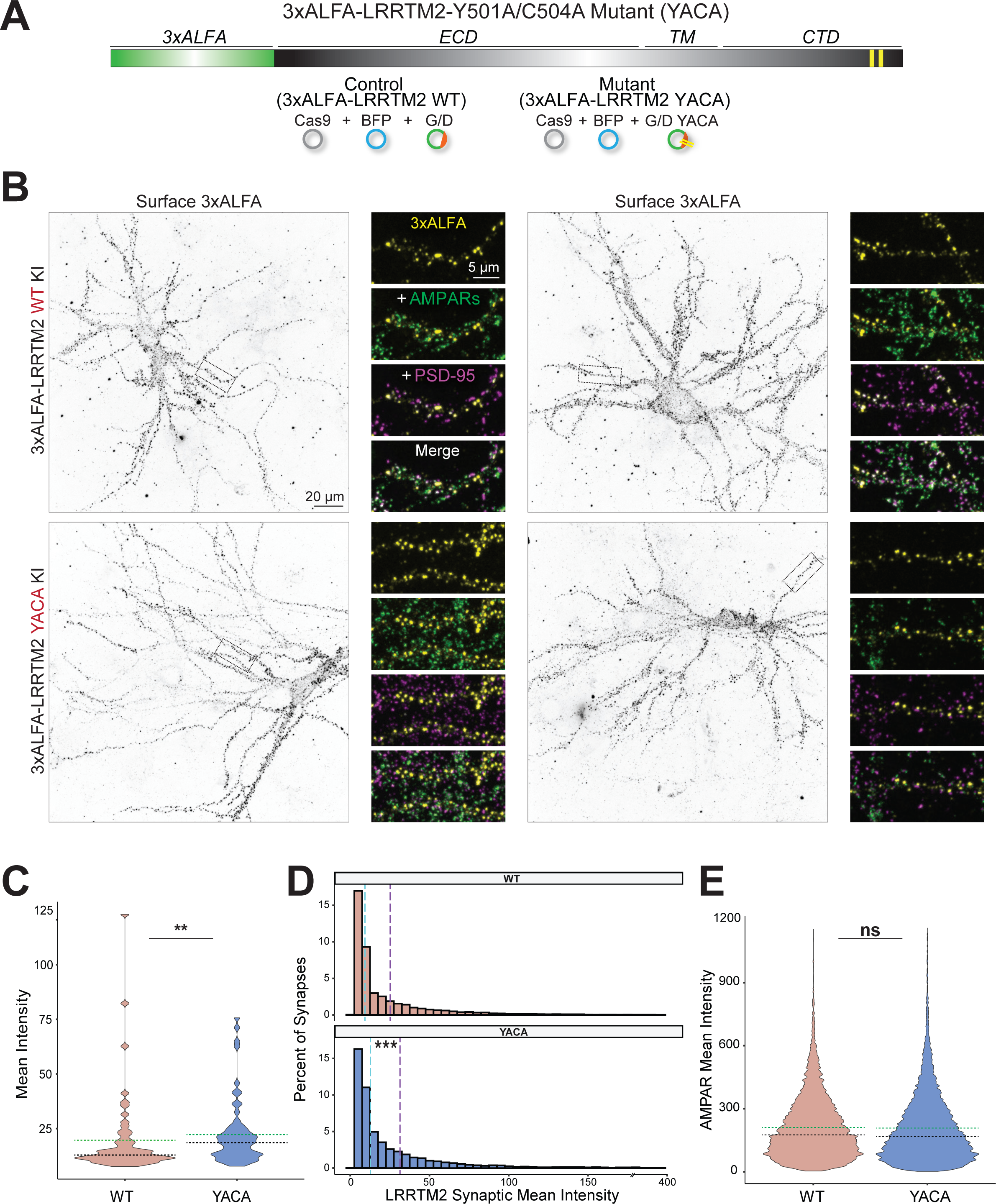
Simultaneous mutagenesis and tagging enabled by whole-CDS TKIT permits analysis of mutation-induced changes in LRRTM2 surface expression **A)** N-terminal 3xALFA tags were inserted in LRRTM2 as in Figure 2, as well as two point mutations in the C-terminus (Y501A + C504A, or YACA). Control condition knock-ins contain tagged wildtype (WT) sequence, YACA condition knock-ins contain both the tag and the two point mutations. **B)** Exemplar images of WT or YACA-mutated LRRTM2 (yellow), co-stained for PSD-95 (magenta) and surface AMPARs (green). **C)** The YACA mutation (navy) of LRRTM2 induces higher synaptic surface intensity than wildtype (tan), averaged by cell (p < 0.005). Green lines represent means and black lines represent medians for each distribution. **D)** Distribution of synaptic enrichment of WT or YACA LRRTM2, showing a rightward shift of the YACA distribution (navy) relative to the WT (tan) (p < 2.2^-16^). Purple lines represent the mean, cyan the median of the respective distributions. **E)** Synaptic surface AMPAR enrichment is not significantly higher in the YACA-mutated condition relative to WT. Green lines represent means and black lines represent medians for each distribution.

Previous studies have established that removing LRRTM2 reduces synaptic AMPAR content^6–8^, and our data indicate that AMPAR content scales with LRRTM2 content at synapses. While we have shown that the YACA mutations moderately increase LRRTM2 content, it is unclear whether this small change would follow the pattern in our correlational data and drive an increase in AMPAR content. We therefore measured synaptic AMPAR staining in each condition and found that the YACA mutations did not have a positive effect on synaptic AMPAR content (Wilcox: p = 1, r-statistic = -0.035; Figure 5E). This suggests that despite the correlation between LRRTM2 synaptic content and AMPAR synaptic content, increasing the LRRTM2 synaptic content by 24.6% via YACA is insufficient or unable to induce a similar increase in AMPAR content.

### Applying whole-CDS replacement to other genes

We conclude that whole-CDS replacement TKIT will be a valuable approach for protein structure-function analysis. The structure of the rat *Lrrtm2* gene, being essentially contained on a single exon, is undoubtedly highly advantageous for the technique, but other genes with different exon/intron structures may still be amenable for the same approach. We presume that the length of the CDS would have to be somewhat modest, but note that the insertion sizes utilized here (3925 bp) are much greater than typically amenable to knock-in via HITI^13^. In fact, the length rather than the content or functional complexity of what is being replaced appears to be most critical for the success of the knock-in. We explored the Santa Cruz Genome Browser for rat genes smaller than a practical limit of 10,000 bases and found 39,459 coding sequences (including splice variants) that had a distance from start to stop codon (CDS spanning region) that is the same or shorter than that of *Lrrtm2* (1882 bp, though the span between our selected guides is 2258 bp) (Figure 6A). Out of the rat genes whose CDS spanning regions are under 10kb, approximately 42.6% are smaller than *Lrrtm2* (Figure 6B). While of course many important genes have long genomic spans, this simple analysis shows that numerous genes may be targetable for whole-CDS replacement. Our conservative estimate reflects the fact that we have not systematically explored the upper range of donor sizes amenable to our approach, suggesting that the actual number of suitable genes may in fact be much higher. Furthermore, while guide selection is undoubtedly critical to knock-in success, the flexibility of guide locations as adapted from TKIT permits more successful knock-in of large donors. Overall, the power of whole-CDS replacement will enable new research on endogenous protein localization, trafficking, and function in neurons avoiding the constraints of overexpression and single-locus CRISPR editing techniques.

**Figure 6:**
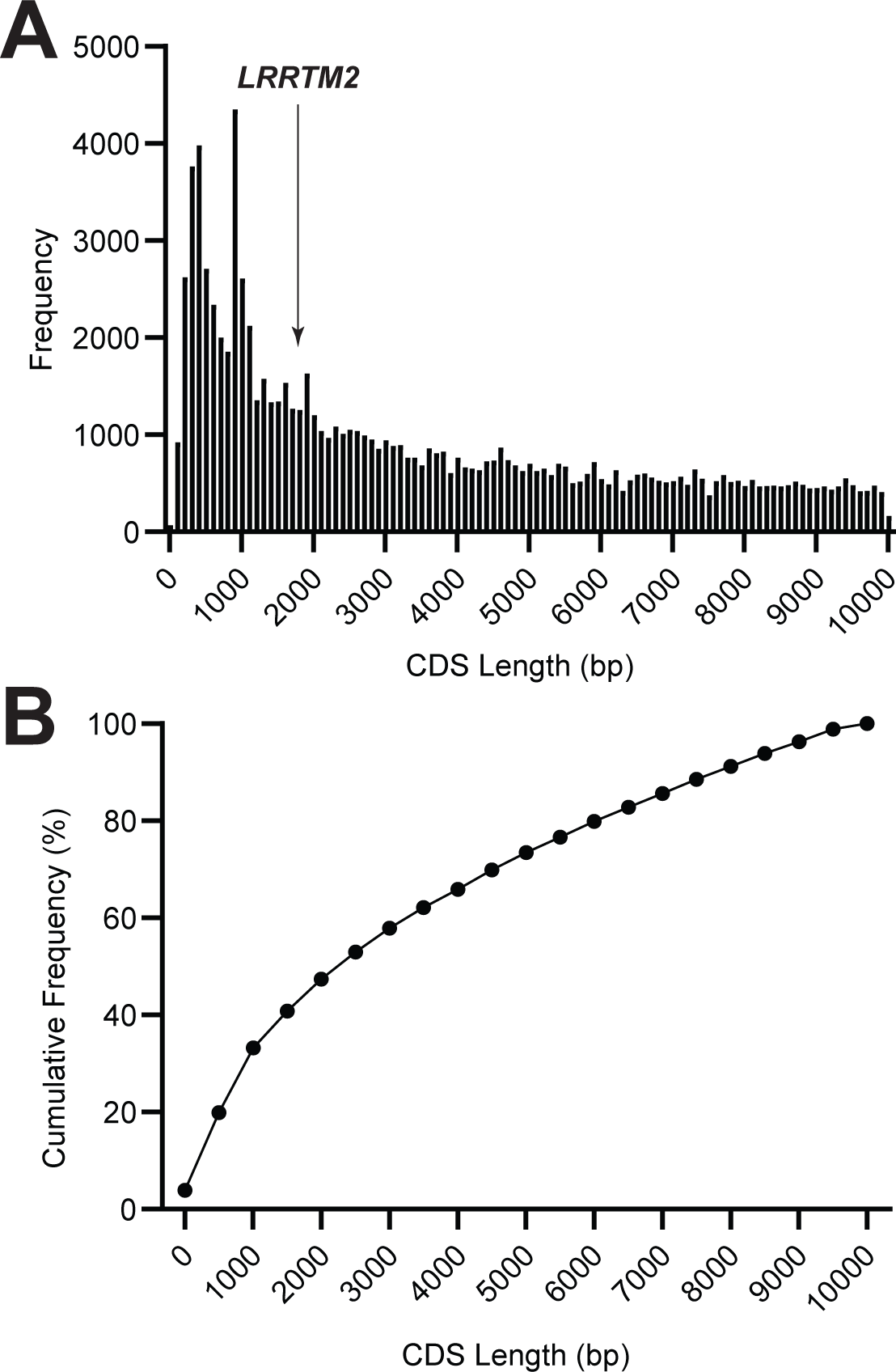
Applying whole-CDS replacement to other genes in *Rattus norvegicus* **A)** Bioinformatics analysis of coding sequence span within the *Rattus norvegicus* genome. Data procured from the Rnor6.0 sequence via UCSC Genome Browser. Results were restricted to complete CDS sequences whose total span across the genome does not exceed 10,000 bases. Arrow shows size of the LRRTM2 CDS, which spans a total of 1882bp from start codon in exon 1 to stop codon in exon 2, with our CRISPR replacement of 2171 bp from guide 1 (intron 1) to guide 2 (3’UTR). **B)** Cumulative frequency distribution of data in A, showing a considerable proportion of the sequences with a CDS spanning region below 10kb are smaller than that of LRRTM2 and therefore potential targets for this method.

## Discussion

In this work, we have demonstrated whole-CDS replacement in neurons, and shown its power to simultaneously tag and mutate a protein at widely separated points of its sequence while maintaining native genetic regulation. The protein-coding portion of thousands of genes span lengths suitable for whole-CDS replacement, suggesting that this simple approach for total control over protein sequence will be a straightforward method for structure-function analysis in diverse systems. We used the approach here to identify new characteristics of expression and trafficking of the critical synaptic adhesion molecule LRRTM2. LRRTM2 levels in cultured hippocampal neurons were unexpectedly variable between neurons, and while its expression correlated with both PSD-95 and AMPAR content, the protein appeared absent from 20% of synapses. We also observed LRRTM2 outside of synapses at puncta that contained AMPARs but lacked PSD-95, suggesting a previously unappreciated role for LRRTM2 outside of PSD-95-containing synapses. Finally, utilizing our ability to manipulate the endogenous LRRTM2 sequence, we were able to increase synaptic LRRTM2 content without affecting AMPAR content, suggesting that this relationship may not be intrinsic or bi-directional.

### Whole-CDS replacement is a general and flexible approach for simultaneous multi-site manipulations

While CDS replacement strategies have been demonstrated in dividing cells^24^, this work is to our knowledge the first demonstration of whole-CDS replacement in neurons. A major advantage of this method is the ability to make multiple, simultaneous modifications to a gene at disparate locations along it in a single editing step. In theory, other CRISPR technologies that allow for DNA replacement could be used for whole-CDS replacement, though likely TKIT, which we relied upon here, is the most generalizable approach. For example, CDS replacement has been achieved in dividing cells for a coding region of similar size to LRRTM2^24^. However, this relied on homology-directed DNA repair, which makes it intractable in post-mitotic cell types such as neurons. This is due to the necessary reliance in post-mitotic cells on Non-Homologous End Joining (NHEJ), a method of donor DNA incorporation that religates blunt ends of DNA together. PRIME editing, which utilizes neither nonhomologous end joining nor homology-directed repair^25^, allows for short substitutions (∼100 bp^26^), but this is too small to replace the vast majority of genes. Designer exon approaches such as CRISPIE^27^ could also be used to essentially “knock-in” an engineered CDS in place of a gene’s first exon and bypassing the later exons, though this would eliminate key intronic regulation and alternative splicing. Finally, while making two separate edits using CRISPR is possible^26,28^, this remains extremely challenging and less efficient than whole-CDS replacement. Further, there is as yet no path for expanding to more than two editing sites, whereas whole-CDS replacement offers near-infinite flexibility within the large donor sequence. Due to the permanent nature of genetic modifications, this whole-CDS replacement approach could simplify the process of generating multiple stem cell lines or animals with multiple genetic mutations in the same gene, as there would be no need to select and validate new guides for each modification that is introduced. Given these factors and the proven efficiency and accuracy of TKIT^12^, we are confident it presents a flexible tool to manipulate entire coding sequences in diverse systems.

The limit on donor size for TKIT is not known, though previous research would suggest that large donor sizes have generally been a struggle for CRISPR in neurons and other models reliant on NHEJ. Previous studies suggest NHEJ may have greater difficulty with larger DNA sizes^29^, though it is also more efficient^13^. It is also conceivable that larger replacements may be facilitated by fusing repair machinery onto Cas9, which has been recently demonstrated to impact its efficiency^30^. Practically, donor size and efficiency likely depend in part on the genetic and epigenetic context as well as practical limits such as plasmid size or viral packaging and must therefore be determined on an individual basis. The former can be addressed with TKIT due to the flexibility of guide locations within the intronic/noncoding regions, which permits higher efficiency guides with lower off-target effects to be selected. We were able to efficiently knock-in a large donor sequence (3925 bp, including IRES-Cre), and while we have demonstrated that a replacement of this size is feasible, replacement of larger regions may certainly also be possible. Furthermore, we have identified a sizeable group of genes whose CDS-spanning region size is smaller than this and thus appear amenable to whole-CDS replacement. Note that for many genes of interest, the difficulty of applying the approach will not be the size of the translated protein but the presence of introns so large as to be impractical to supply in a donor. In such cases, it may still be possible to adapt this technique to replace regions on one side or the other of a large intron, including multiple small, contiguous exons and introns, without modifying the entire CDS. This approach would largely preserve the regulation intrinsic to intronic sequences such as transcription and alternative splicing.

### Whole-CDS replacement permits versatile tagging and reporter strategies

Ready identification of knock-in cells is key to experimental performance in many techniques, including microscopy and flow cytometry. However, knocked-in tags can be difficult to visualize because endogenous protein expression is often far lower than typical overexpression, and reliance on abundant target protein translation for tag detection is a potential source of bias. While empirical determination of the brightest protein tags can help improve utility, a knock-in conditional marker independent of the target protein’s expression level is highly useful, and essential in many cases. By integrating existing Cre-FLEx conditional marker technologies into non-coding regions simultaneously with our modifications to the LRRTM2 coding sequence, we were able to identify, tag, and manipulate knock-in cells in a single step. Importantly, this strategy has previously only been available for C-terminal knock-ins, or N-terminal using a Cre-P2A sequence^31,32^, whereas whole-CDS replacement allows combining this manipulation with other mutations throughout the CDS. The FLEx system is highly adaptable, and the cell fills we utilized here can be exchanged for any genetically-encoded protein marker, sensor, or optogenetic tool desired. This combination of powerful tools in an efficient CRISPR environment should permit elegant, high-throughput studies of protein function in their cellular context in the future.

### Endogenous LRRTM2 expression varies across and within neurons

One surprising result from this study was the cellular variability of LRRTM2 expression even within the relatively restricted range of neuron types found in hippocampal culture. LRRTM2 mRNA levels have been found to vary between cell types in neocortex and hippocampus^33,34^. The correlation in single cells between mRNA and protein levels is frequently poor, yet analysis of population protein expression levels at single-cell resolution is challenging. Here, sparse knock-in allowed systematic evaluation of single-neuron expression as well as the measurement of expression levels of mutant protein. Note that such analysis is aided substantially by the presence of a knock-in dependent marker, which we expect will be particularly useful for analysis *in vivo* where the range of expression levels may be even greater. The mechanisms of cell-specific LRRTM2 expression level remain unknown, but one attractive possibility is that the activity history of the neuron drives regulation of LRRTM2 as the number of AMPARs and synapses is modulated.

Our data have shown conclusively that LRRTM2 is present at higher levels in larger synapses. LRRTM2 binds directly with PSD-95 via its PDZ-binding motif^4,10^, and endogenous LRRTM2 was indeed most abundant at synapses with more PSD-95 content. Furthermore, we were able to increase the synaptic content of LRRTM2 via the YACA mutations. Notably, this increase in synaptic content was smaller than expected from mutant overexpression^22^, highlighting the importance of manipulating LRRTM2 trafficking, and protein trafficking more generally, with endogenous regulations intact. While the mechanisms by which the YACA mutations influence endogenous LRRTM2 trafficking are unclear, the intracellular location of the YACA mutations suggests intracellular interactions may be involved in trafficking LRRTM2, including possibly PSD-95 whose interaction domain is close by. However, we also found that LRRTM2 was not present at nearly 20% of PSD-95-containing synapses and was also present at dendritic locations lacking PSD-95, suggesting there are additional mechanisms beyond binding to PSD-95 responsible for LRRTM2 localization. Several overexpression studies have found that removing the PDZ-binding motif does not affect the ability of LRRTM2 to traffic to synapses or facilitate LTP^7,10^, suggesting it is not exclusively responsible for LRRTM2 trafficking. Furthermore, it appears that these potentially extrasynaptic LRRTM2 puncta frequently contained AMPARs. What would prompt enrichment of LRRTM2 at these points is unclear. The neurexin-binding domain of LRRTM2 is required for rescuing the effects of removing LRRTM2 on LTP via overexpression^6,7^, but it is unclear whether neurexins, or indeed any presynaptic scaffold, would be present outside of PSD-95 synapses. One hypothesis is that these LRRTM2 puncta are indeed synapses but lack PSD-95 and contain other synaptic scaffolds whose interaction with LRRTM2 has not been investigated. Given the homology in PDZ motifs across synaptic scaffolds, it is certainly likely that the LRRTM2 PDZ-binding motif also interacts with other scaffolds such as SAP-102 or PSD-93^35^. However, previous research has found that SAP-102-only synapses are relatively rare past early postnatal development^36,37^, and as such are unlikely to account for very many of these LRRTM2 puncta. Alternatively, it is possible that these puncta represent a diffusible pool of LRRTM2 that can be recruited to the synapse as needed, much the way that AMPARs are^20,21^. This model would include two pools of LRRTM2 protein, one synaptic and one extrasynaptic, where the synaptic pool would be either non-functional or diminished in size when unable to bind presynaptic neurexins but not PSD-95 or other scaffolds.

LRRTM2 and its sister protein LRRTM1 share high sequence homology and are expressed in the same broad regions of hippocampus^38^, and many studies have relied on their simultaneous deletion^6,7^. This is done because CAMs frequently compensate for one another, and dual knockouts can better illustrate their roles than either knockout alone, as has been recently shown for CAMs LRRTM1 and SynCAM1 in hippocampal synapses^39^. Interestingly, the effects of dual knockout can be rescued with overexpression of LRRTM2 alone^6,7^. We found endogenous LRRTM2 in 80% of synapses, which raises the possibility that the remaining 20% may be LRRTM1-positive. This is potentially an interesting case of differential subcellular trafficking of highly homologous proteins. A more extreme segregation between members of a single family of CAMs has been observed previously *in vivo*; for example the neuregulin proteins 1 and 3 are differentially sorted to the somatic and axonal domains of pyramidal neurons, respectively^40^. The functional implications of such segregation are unknown at this point, but given that many studies have relied on their simultaneous deletion, future work is needed to tease out their individual roles. While it is currently extremely challenging to simultaneously target multiple genes via CRISPR, future developments could make it possible to tag and manipulate both LRRTM1 and LRRTM2 in the same cells to further investigate their relative trafficking.

### Relationship of endogenous LRRTM2 with AMPARs

Numerous previous studies have established that LRRTM2 controls synaptic AMPAR content. One possibility is that changing LRRTM2 levels would be determinative in establishing AMPAR levels. Our data show that synapses lacking LRRTM2 are smaller and contain fewer AMPARs than those with LRRTM2, though the latter correlation was weaker than predicted by this hypothesis. However, when we increased the content of LRRTM2 with the YACA mutations, there did not appear to be a similar increase in synaptic AMPAR content. Several possible explanations exist for this finding. One possibility is that while YACA mutations may increase LRRTM2 surface content by altering membrane-targeting trafficking mechanisms^23^, they also block the as yet unknown mechanisms by which LRRTM2 anchors AMPARs at the surface. However, given that previous studies have removed the entire C-terminal domain of LRRTM2 in a replacement context with no effect on LTP^6,7^, this appears unlikely. Another possibility is that the YACA mutations, which lead to a cell-wide increase in LRRTM2 content, may not significantly alter AMPAR trafficking at the cellular level due to potential compensatory mechanisms. Instead, it’s plausible that AMPAR trafficking could be influenced by localized increases in LRRTM2 content on a synapse-by-synapse basis. Further, it is possible that the mechanism connecting AMPAR enrichment to LRRTM2 levels is not sensitive enough to respond to the small increase that the YACA mutations induce in LRRTM2 content. Unfortunately, these latter two possibilities are difficult to test without a greater understanding of what induces LRRTM2 trafficking to synapses so that it can be manipulated more directly and in a physiologically relevant manner. Finally, it is important to note that we measured the LRRTM2 YACA mutation at a static baseline state, whereas it is possible that the role of the domain containing the YACA mutants is engaged primarily during synaptic potentiation.

Previous studies have established that surface diffusion of AMPARs to synapses is critical during plasticity^20,21^. Our data have shown that LRRTM2 and AMPARs localize to puncta outside of PSD-95 and appear together more frequently than either protein alone. This suggests the exciting possibility that LRRTM2 may traffic together with, or potentially shepherd, extrasynaptic AMPARs as they are trafficked to synapses during plasticity. This would give LRRTM2 the ability to retain new AMPARs at synapses via its interaction with presynaptic neurexins in a timely fashion. If LRRTM2 recruits AMPARs to the synapse via this surface trafficking mechanism, it would explain why a whole-cell increase in LRRTM2 content, as induced via the YACA mutation, is insufficient to induce a similar increase in synaptic AMPAR content. LRRTM2 facilitates synaptogenesis^4^, and extrasynaptic LRRTM2-AMPAR puncta suggest a means by which LRRTM2 could enable AMPAR trafficking to new synapses. The presence of LRRTM2 clusters outside of PSD-95 synapses, and their potential role in local AMPAR trafficking, represents an intriguing new development in our understanding of LRRTM2 function and requires further investigation to fully understand.

Together, our approach for whole-CDS replacement facilitates labeling, imaging, and manipulating endogenous proteins, including LRRTM2, and represents a potent methodological advance in the field of cellular neurobiology. While we have illustrated many new findings regarding the endogenous trafficking of LRRTM2, the approach of whole-CDS replacement will assist in determining the mechanisms by which this cell adhesion molecule controls such critical processes as synaptic AMPAR retention and plasticity.

## Methods

### CRISPR Design and Plasmids

The NCBI Rnor 6.0 database LRRTM2 sequence (NC_005117.4) was used as a reference for both guide and donor design. Guides positioned at least 50bp from the intron-exon splicing boundary (5’ guide) or over 50bp into the 3’UTR (3’ guide) were identified using the Benchling CRISPR guide ranking tool, targeting those with the best on-target and off-target scores based on previous publications^41,42^, then were cloned into the pX330 backbone as previously described^12^. Guide sequences: 5’ GTTTTAATCTCTCTTATACA 3’ (guide 1, anneals to Intron 1) and 5’ CTTTTTAAGTAGGAAGCCAG 3’ (guide 2, anneals to the 3’UTR). Once cloned into the pX330 vector under identical U6 promoters, the guide construct was grown in NEB STABLE cells at a reduced temperature of 30°C to prevent bacterial recombination of the promoters. Guide 2 was cloned with a guanine at the 5’ end after the U6 promoter, as described in Fang et al. 2021. 3xALFA epitope tag and IRES2-Cre insertions as well as YACA (Y501A/C504A) mutations were added to the donor by either Gibson NEB HIFI Assembly or NEB Q5 site-directed mutagenesis as appropriate. The 3xALFA tag was inserted after the signal peptide along with a small linker (TS) after the tag, and IRES2-Cre was added immediately after the stop codon. For lentiviral expression, the guides and donor were combined into the pFW lentiviral backbone^18^ by NEB HIFI Assembly. FLEx-mTagBFP2 was generated by NEB HIFI Assembly, replacing the mCherry-KASH in Addgene #139652 (a gift from Harold MacGillavry; http://n2t.net/addgene:139652; RRID:Addgene_139652) with the mTagBFP2 gene and replacing the hSyn promoter with the CAG promoter for improved expression levels. FLEx-IRES-EGFP was made similarly, with a CMV promoter in place of CAG. HA-spCas9 was a gift from Harold MacGillavry (Addgene plasmid # 131506; http://n2t.net/addgene:131506; RRID:Addgene_131506), constructs to make lentivirus (pMD2.G and psPAX2) were gifts of Didier Trono (Addgene plasmid # 12259; http://n2t.net/addgene:12259; RRID:Addgene_12259; Addgene plasmid # 12260; http://n2t.net/addgene:12260; RRID:Addgene_12260). ALFA and HA tagged LRRTM2 KDR variants for transient expression in HEK cells were generated by replacing the GFP tag in KDR GFP-LRRTM2 previously described^8^ using NEB HIFI Gibson cloning. All sequences were confirmed by whole plasmid sequencing (Plasmidsaurus) using Oxford Nanopore Technology with custom analysis and annotation. These plasmid sequences will be deposited with public databases upon publication.

Note that we originally tried to combine the FLEx fluorescent protein with the guides/donor lentiviral plasmid, but full plasmid sequencing revealed that growing a construct with both IRES2-Cre and FLEx genes in bacteria caused a reversal of the flipped gene. Based on previous literature, we suspect that this could be due to bacteria recognizing the IRES2 sequence and expressing the Cre protein themselves^43^. Thus, we elected to separate these constructs into two viruses, with the Cas9 virus as a third virus due to its large size and the limitations of lentiviral packaging. The guides and Cas9 were cloned into separate viral constructs to avoid Cas9 activity at the *Lrrtm2* locus in the absence of donor sequence to replace it.

### Genomic sequencing

Cultured hippocampal neurons were infected with either knock-in viruses (Cas9 + guides/donor) or guides/donor alone at DIV1. DNA was extracted and purified at DIV21 using the Wizard Genomic DNA Purification Kit (Promega) according to manufacturer’s instructions. The LRRTM2 gene was amplified using primers that anneal on either side of the guide1 Cas9 cut sites (5’ AGCCAGTGAATTCCCGTTTT 3’, 5’ AGGCGAACTGGGATAGTCCGCA 3’). This PCR product was gel-purified, then PCR amplified again using a reverse primer that anneals specifically to the knock-in HA sequence (5’ CATTTAGGTGGACAACTAGTAGCGTAGTCTGGTACATCAT 3’) and the same forward primer as in the first PCR run. Gel purified PCR products from the round 1 PCR control condition and the round 2 PCR knock-in condition (no product was made from the control condition due to the lack of knock-in HA sequence in the *Lrrtm2* locus) were Sanger sequenced (Genewiz/Azenta) to confirm knock-in in the correct genetic locus.

### Lentivirus Production

HEK293T cells (ATCC CRL-3216) maintained in DMEM + 10% FBS supplemented with penicillin/streptomycin were plated at high density and transfected using PEI as we have described^18^. Cells were incubated for 6 hours followed by a media change to standard neuronal culture media. Lentivirus was allowed to accumulate in the media for 2 days before harvesting, and debris were removed by centrifugation at 1000 x RPM for 5 minutes. Virus aliquots were used fresh or else frozen at -80°C. Based on prior titrations and to achieve high levels of co-infection, neurons were infected with 100μL of each virus.

### Immunostaining

Dissociated, mixed-sex hippocampal cultures were prepared as previously described^18^. Knock-in preparations were infected at DIV1 with 100 ul each unconcentrated Cas9, guide/donor, and marker viruses. To immunostain LRRTM2 and synaptic proteins, coverslips were removed from their culture media at DIV21 and blocked for 5 minutes in 1% BSA in Tyrode’s buffer, then live-labeled with ALFA-AlexaFluor647 nanobody (NanoTag) and mouse anti-pan GluA (Synaptic Systems) at 1:500 and 1:200, respectively, in the same solution composition for 12 minutes at room temperature. To avoid background from aggregation, aliquots of nanobody are vortexed thoroughly for 2 minutes followed by centrifugation at max speed for 1 minute. Coverslips were then rinsed twice briefly in phosphate buffered saline (PBS) and fixed in 4% paraformaldehyde (PFA) + 4% sucrose in PBS for 9 minutes at room temperature. The cells were washed with PBS-100mM Glycine (PBS-Glycine) three times for five minutes each, permeabilized with 0.3% TritonX-100 in PBS-Glycine for 15 minutes at room temperature, then incubated with PSD-95 nanobody conjugated to AZ568 dye (NanoTag) at 1:100 and goat anti-rabbit AlexaFluor488 at 1:750 in 4% TBS-milk for 45 minutes at room temperature. Coverslips were then rinsed in TBS three times for five minutes prior to imaging.

To test different knock-in tags, we transfected HEK cells using Lipofectamine 2000 (ThermoFisher) with an EGFP cell fill to normalize for transfection as well as the respective KDR LRRTM2 constructs with 1xALFA, 3xALFA with interstitial linkers, and a 3xALFA without linkers. A 1xHA tag visualized with a traditional primary-secondary antibody approach was included for comparison. We then immunolabeled, fixed, and stained as above. Cells were fixed and surface stained with either the ALFA nanobody or an HA primary antibody followed by fluorescent secondary. Both approaches utilized the AlexaFluor 647 dye to detect LRRTM2 as well as identical imaging parameters. Images were analyzed in ImageJ for 647 fluorescence intensity normalized to GFP as a transfection control.

### Microscopy

Images were acquired on an Andor Dragonfly spinning disc confocal (Andor) attached to a Nikon Ti2 Eclipse inverted microscope base with a 60x Plan Apo 1.49 NA objective. Excitation laser light (405, 488, 561, or 638 nm) from an Andor ILE, flattened by an Andor Beam Conditioning Unit, was passed to the sample by a 405/488/561/640 quadband polychroic (Chroma). Emission light was passed through an appropriate bandpass filter (FF02-447/60-25 (Semrock), ET525/50, ET600/50 (Chroma), or Em01-R442/647 (Semrock), for 405nm, 488nm, 561nm, and 638nm emission, respectively) and collected on a Zyla4.2 sCMOS camera. Cells of interest were imaged with confocal z-stacks with 0.5 μm z-steps at 25-90% laser power with 200 ms exposure (400 ms for 638nm channel), with each channel imaged sequentially.

### Analysis

Image processing was performed using macros and plugins in Fiji/ImageJ^44^, and image file names were blinded during analysis. All z-stacks were converted to maximum intensity projections, then *xy* chromatic aberrations were corrected using the *Register Channels* tool of the NanoJ – Core plugin^45^ (and 4-color TetraSpek bead images acquired prior to imaging as a standard.

We used a semi-automated ImageJ macro similar to that described in Dharmasri et al^18^ to detect synapses. In brief, the macro provides user-guided image cropping, followed by automated thresholding to isolate ALFA-LRRTM2 signal, then uses the puncta detection plugin SynQuant^19^ to detect synapses on knock-in neurons from the PSD-95 staining. Detected synapses were converted to ROIs and used to measure synapse area and PSD-95 intensity, as well as intensity of LRRTM2 and/or AMPARs within those puncta. Due to high cellular variability, we took the cellular average for each channel and assigned each synapse ROI a z-score based on their relative intensity within that channel to the cellular average.

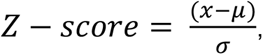

where *x* is the intensity at the individual synapse, *μ* represents the mean synaptic intensity across the neuron, and *σ* represents the standard deviation. This was performed for two measures of intensity readout: Mean Intensity (Figure 3C) and Raw Integrated Density (everything else), the latter of which is superior for establishing protein content correlations as it does not normalize for synapse size, which is biologically relevant. We observed infrequent outliers on the upper end of the intensity distributions across all three staining conditions (LRRTM2, PSD-95, and AMPARs), presumably due to clumping of the antibody or nanobody. To eliminate these synapses from our dataset in a way that was agnostic to the normality of the distribution, we utilized a cutoff of 150% of the interquartile range (IQR) above the third quartile (Q3 + 1.5*IQR) in each of the three channels’ distributions. These outlier calculations were performed using the z-scored data to avoid artificially removing synapses from cells with high expression levels of any of the measured proteins, particularly LRRTM2. Synapses above the calculated cutoff were removed completely from the datasets.

Synapses along knock-in dendrites were categorized as containing or lacking LRRTM2 by comparing the distributions of ALFA labeling along knock-in dendrites or neighboring dendrites (acquired simultaneously and analyzed with the same pipeline as above in parallel).

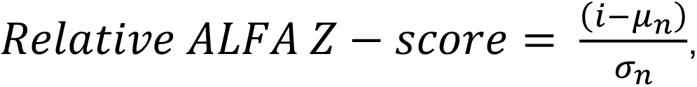

where *i* is the synaptic ALFA intensity (KI dendrite or neighboring region), *μ_n_* represents the mean synaptic ALFA intensity of the adjacent neighboring region, and *σ_n_* represents the standard deviation of the adjacent neighboring region. The 95^th^ percentile of the ALFA intensity distribution on neighboring dendrites was used as the cutoff between “LRRTM2-lacking” and “LRRTM2-containing” synapses along knock-in dendrites.

For detection of LRRTM2 and/or AMPAR-containing puncta within and outside PSD-95 puncta, we hand-picked elliptical ROIs around puncta in each channel using FIJI ROI selection tools, then quantified fluorescence intensities as above. Raw Integrated Density was measured within each hand-drawn ROI, followed by background subtraction using similarly-sized ROIs drawn nearby in the background region of each image. Each puncta was categorized as “positive” or “negative” for each of LRRTM2, PSD-95, and AMPARs utilizing a first local-minimum cutoff in the intensity distribution of each protein across the dataset. These categories were plotted as pie charts using a combination of RStudio and GraphPad Prism.

### Bioinformatics

To assess how applicable a whole-CDS replacement of a similar size to our approach would be to other genes, we turned to a bioinformatics approach. Genetic data on *Rattus norvegicus* was pulled from the search function of the University of California Santa Cruz Genome Database (https://genome.ucsc.edu/cgi-bin/hgGateway). To measure the CDS Spanning Region along the genome for each gene and splice isoform, we subtracted the stop codon position from the start codon position. Practically, replacement size will depend on the proximity of suitable PAM sites and guide sequences on either side of the CDS. For practicality of graphing, we set the maximum CDS Spanning Region size as 10,000bp. Search parameters entered into the UCSCGD were as follows: absolute value(endCDS – startCDS) <= 10000bp. Alternative splice isoforms of the same gene were included as separate genes because they likely differ in genomic span and carry differing biological significance. Histogram was produced in RStudio, cumulative plot in GraphPad Prism.

### Statistics

Data from ImageJ measurements was processed and statistics calculated using RStudio (Posit). Frequency distributions were produced for each of the synaptic intensity measurements to evaluate normality. These distributions were not normally distributed as evaluated with a Shapiro-Wilks test, and we therefore elected to utilize Wilcoxon tests to compare LRRTM2-containing and -lacking synapses, as well as YACA and wildtype LRRTM2 mutant conditions. Graphs were produced in RStudio or GraphPad Prism (Dotmatics). Due to the high throughput nature of our semi-automated synapse detection method, we pruned the number of points displayed in scatter plots by averaging every 5 synapses together based on ranked data (PruneRows, Prism). While this had very little effect on the slope of the regression lines, it does influence the R-squared values so we have reported the values of regressions performed on the raw data in the text.

## Contributions

The project was conceptualized by SLP and experiments designed by SLP and TAB. TKIT CRISPR constructs were designed and created by SLP and ADL. Experimental conditions and reagents were optimized by SLP and MCA. Data were collected and analyzed by SLP. TAB supervised the study. Figures were prepared and the manuscript written by SLP with constructive review and editing by TAB, MCA, and ADL.

## Funding

SLP – F32MH130106

MCA – F31MH12428330

TAB – R37MH080046, R01MH119826

