## Supplemental Figure 1 for "Large donor CRISPR for whole-CDS replacement of cell adhesion molecule LRRTM2" (from another version of this preprint: version 1, 2024-05-05)

[3] Program in Neuroscience, University of Maryland School of Medicine, Baltimore, MD, USA
[*]corresponding author

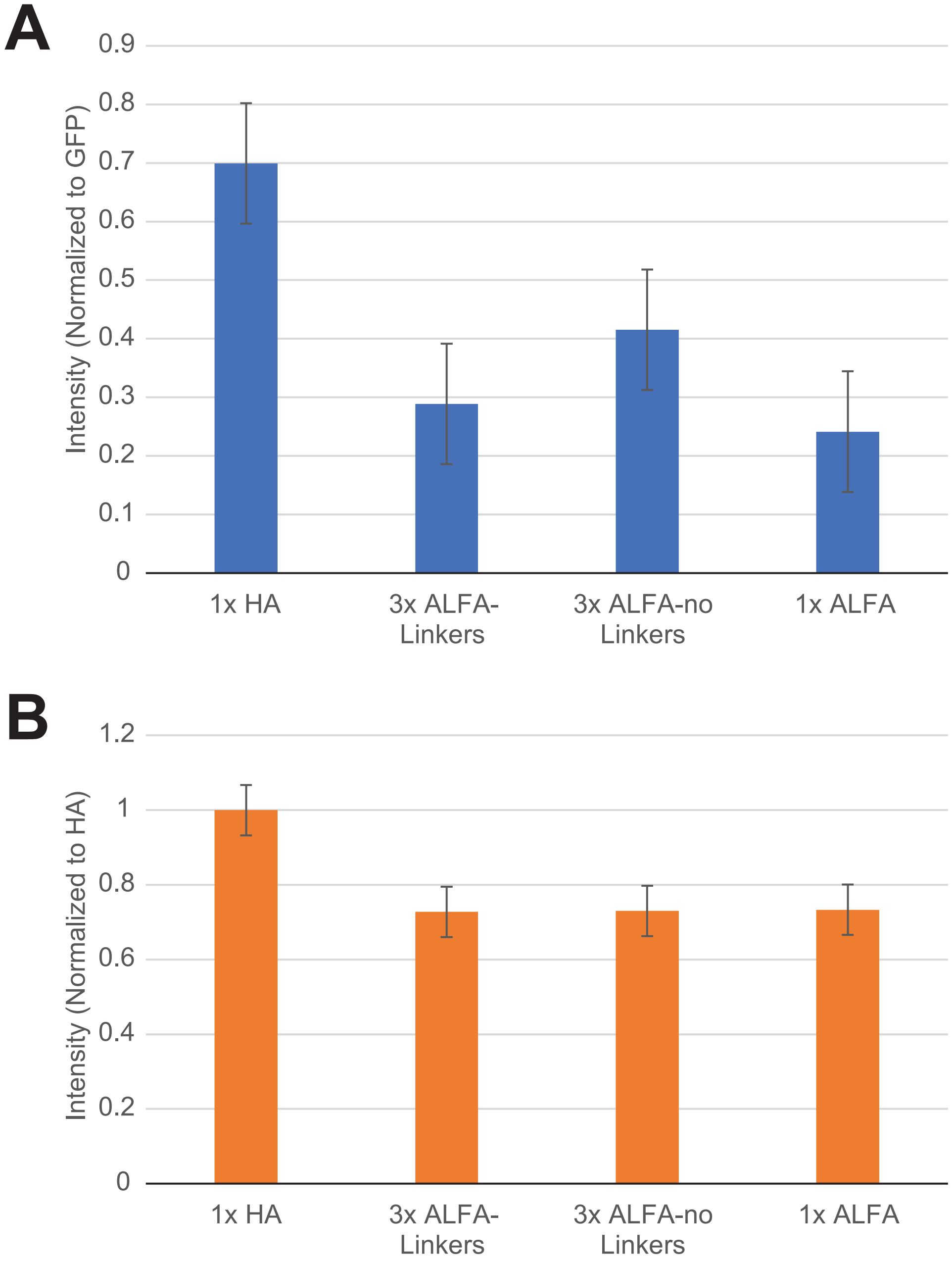
A
0.9
0.8
0.7
0.6
0.5
0.4
0.3
0.2
0.1
0
Intensity (Normalized to GFP)
1x HA
3x ALFA-Linkers
3x ALFA-no Linkers
1x ALFA
B
1.2
1
0.8
0.6
0.4
0.2
0
Intensity (Normalized to HA)
1x HA
3x ALFA-Linkers
3x ALFA-no Linkers
1x ALFA

**Supplemental Figure 1**: Triple ALFA tag without interstitial linkers leverages low background of nanobody staining with superior amplification to a singular tag

**A)** To determine the best tag for visualizing endogenous LRRTM2, we evaluated three versions of the epitope tag ALFA, which can be stained using a dye-conjugated nanobody: 1xALFA, 3xALFA with interstitial linkers, and a 3xALFA without linkers. A 1xHA tag visualized with a traditional primary-secondary antibody approach was included for comparison. Measured intensities were normalized for transfection using co-infected GFP cell fill. **B)** Background fluorescence intensity analyzed across conditions from A, normalized to the HA condition. While the primary-secondary antibody amplification provides a higher signal (**A**), secondary antibody-induced background fluorescence is also higher (**B**). Interestingly, background ALFA levels are low, and tripling the ALFA tag improves signal by approximately 50% but only when no interstitial linkers are included.
